# Growth and molecular responses of potato to lunar regolith simulants

**DOI:** 10.64898/2026.02.23.707481

**Authors:** David Handy, Anika Loeffler, Medora Knudson, Sydney Campbell, Pankaj Jaiswal, Jeffrey C. Anderson, Aymeric Goyer

## Abstract

**Background:** On-site food production will be required to achieve NASA’s goal of a sustainable Lunar habitat. Toward this end, the use of fine, soil-like material on the Lunar surface, known as regolith, has been proposed as a plant growth substrate. However, how this substrate may affect plant growth is not well understood. Lunar regolith is devoid of the organic materials that make soils on earth fertile for plant growth, and has been weathered by solar winds, cosmic rays, and micrometeorite impacts. Additionally, regolith at certain lunar sites may contain heavy metals. These metal ions may leech, thus posing challenges with accumulation in plant material. To address and verify the efficacy of regolith-based crop production, we used lunar regolith simulants (LRS). We investigated the effects of LRS on potato (*Solanum tuberosum* cv ‘Modoc’) plant and tuber development, gene expression, and nutrition profiles.

**Results:** Growth in LRS negatively impacted the potato plant size and tuber yield. While the degree of impact differed between simulants, all plants grown in LRS were statistically significantly shorter in height than plants grown in control soil. Further experiments with the lunar mare simulant 1E (LMS-1E) show that these effects can be ameliorated through the addition of vermicompost, an organic component, with a 70:30 v/v ratio of LMS-1E to compost being virtually indistinguishable from controls. Changes in gene expression profiles also differed between simulants, with genes related to photosynthesis, biotic and abiotic stress responses, signaling, and terpenes and flavonoids metabolism being commonly altered. Despite these observed differences in transcription, broad changes in metabolite profiles were not observed.

**Conclusions:** LRS are clearly stressful on plants. However, amendment of the substrate with composted materials appears to be a viable strategy to alleviate stress. Given these observations, regolith-based agriculture may not be viable for very early food production when organic matter content is low. However, this would improve over time with continual incorporation of organic matter to regolith. As such, we believe regolith-based agriculture is a viable long-term strategy.

## Background

It is the goal of NASA’s Artemis missions to establish a permanent human presence on the Moon in the 2030s (https://www.nasa.gov/feature/artemis/). However, achieving a sustainable and affordable presence on the lunar surface will require on-site food production to reduce resupply costs and increase mission success probabilities [1]. Regolith-based agriculture has been proposed as an *in-situ* method of supporting crops for food production. However, surface lunar regolith as a viable substrate needs to be verified to ensure adequate food production and food safety. As genuine lunar regolith is not available for large-scale plant studies, lunar regolith simulants (LRS) are commonly used for experimentation [2–4].

Lunar regolith falls into two categories: lunar mare and lunar highlands. The primary component in lunar mare regolith is basalt formed by lava flows during the volcanically active phase, while the geologically older, crater-filled lunar highlands primarily consist of ferroan anorthosite [5]. The most unique feature of lunar regolith compared to terrestrial soils and LRS is the presence of agglutinates, aggregate particles formed by micrometeorite impacts and comprised of nanophase iron, mineral fragments, and glass. While LRS often appropriately contains volcanic glass, agglutinates are more difficult to manufacture and are often excluded from LRS. While lacking nitrogen, lunar regolith and its simulants contain micronutrients such as phosphorous, potassium, magnesium, iron, manganese, and calcium [6,7]. While these nutrients may appear replete, it should be noted that the bioavailability of some, such as phosphorous, is dependent on pH and the presence of other elements.

Potato (*Solanum tuberosum*) is an ideal candidate for space crop production because it is high-yielding, clonally propagated, rich in calories and phytonutritients, and has a high satiety index. Potato is also a good model organism for plants that produce underground tubers. It has a rich history of available genetic resources in the form of germplasm accessions, sequenced genome and public access to many large-scale omics datasets for undertaking comparative studies. Early experiments involving lunar samples returned from Apollo-era missions exposed potato seedlings, among other plants, to suspensions of lunar regolith with no reported effects on plant development [8]. Most recently, *Arabidopsis thaliana* grown in lunar regolith returned from three Apollo sites had stunted roots, and slowed aerial growth compared to a LRS, and each regolith induced unique overall gene expression changes [9]. However, plants showed abbreviated growth and arrest in the vegetative stages, activated stress response pathways and negatively impacted the telomerase activity and shortened telomeres[10]. Negative effects on potato tuber yield has been reported when grown in Martian regolith simulant [11], but the effects of LRS on potato growth and yield has not yet been ascertained.

Here, we present findings from two greenhouse experiments performed on the potato cultivar ‘Modoc’. In the first experiment, we tested potato growth on LRS amended with different percentages of organic matter. This enabled us to find a ratio LRS/compost that supports sufficient plant growth for analyzing growth phenotype traits, physiology, and metabolism but is also suboptimal to investigate the stress response to LRS. Data was used to design our second experiment. In the second experiment, we assessed the effects of a variety of LRS types mixed with compost on growth, physiology, leaf and tuber metabolite profile, and leaf gene expression. To our knowledge, this is the first report of the response of potato to LRS with fully-grown plants.

## Methods

### Chemicals

Chlorogenic acid, HPLC-grade acetonitrile, HPLC-grade methanol, and meta-phosphoric acid were from VWR (Radnor, PA, USA). L-Phenylalanine, L-tyrosine, ascorbic acid, solanine, chaconine, methoxamine, pyridine, and ribitol were from Millipore-Sigma (St. Louis, MO, USA). N-methyl-N-(trimethylsilyl) trifluoroacetamide with chlorotrimethylsilane was purchased from CovaChem (Loves Park, IL, USA).

### Potato and growth substrate materials

Mini-tubers of the potato cultivar Modoc were obtained from CSS Farms (Colorado City, CO, USA). Lunar regolith simulants LHS-1E and LMS-1E were purchased from the Exolith Lab (Orland, FL, USA), and JSC-1A, OPRH4W30, and NUW-LHT-5M were obtained from NASA. The Adkins-series soil was obtained from the top 30-cm layer of a non-cultivated, Rye grass field from Umatilla County, Oregon (45.817318° N - 119.294922° W). Vermicompost was sourced from Earthworm Castings (Green Bay, WI, USA).

### Growth conditions

#### Experiment 1: Determination of optimal LRS/compost ratio

Vermicompost was selected as the green compost to mimic the use of inedible plant matter as a soil amendment in a bioregenerative life support system. Mixtures were prepared by combining 5% non-autoclaved vermicompost with 10% or 25% autoclaved vermicompost in 85% or 70% LMS-1E (volume ratios), respectively. The consistent amount of non-autoclaved vermicompost between batches enabled us to obtain mixtures with the same starting microbiome. Adkins-series soil was used as a control of optimal potato growth and was mixed with vermicompost in the same ratios as LRSs. Mini-tubers were planted on 2 August 2024 in 8.89 x 7.62 cm square pots lined with landscaping fabric (Greenhouse Megastore, Danville, IL, USA), containing approximately 300 cm^3^ of soil mixture. Each treatment was done with four biological replicates with one plant per pot. Pots were placed in a fixed, randomized configuration under a PlantEye600 scanner (Phenospex, Heerlen, The Netherlands) for phenotyping, with a border row of plants around the experimental plants to reduce edge-effects. Plants were grown using natural sunlight in a greenhouse set to 22℃ day and 18℃ night. Plants were watered with 40 mL of reverse osmosis filtered water every 2-3 days, increasing frequency as plants matured.

Plants were scanned using the PlantEye600 scanner every 12 hours for the first six weeks after emergence until plant foliage from different plants overlapped (Table S1 & S2). The scanner recorded digital biomass, greenness, height, hue, leaf angle, leaf area, normalized difference vegetation index (NDVI), normalized pigments chlorophyll ratio index (NPCI), and plant senescence reflectance index (PSRI). At nine weeks, a MultispeQ (PhotosynQ Inc., East Lansing, MI, USA) was used to analyze plant leaves for pigment content, and photosynthetic parameters such as quantum yield, electrochromic shift, non-photochemical quenching, etc. (Table S3 &S4). Plants were harvested on 15 October 2024, 74 days after planting (DAP). Plants were removed from the substrate and the roots and tubers rinsed with tap water. The substrate was sieved to remove any remaining roots, stolons, and Osmocote pellets, then collected for chemical and microbial analysis (see below). For additional phenotyping measurements, total mass of above- and below-ground tissues was first measured, then plant parts were separated to measure shoot (leaves and stems) mass, root mass, and tuber mass. Tuber and total masses were used to calculate harvest index as a percentage of edible biomass (100 x edible biomass/total biomass). Leaves, roots, and tubers were flash frozen in liquid nitrogen, lyophilized, and ground to powder with mortar and pestle for further analysis.

#### Experiment 2: Effect of various lunar regolith simulant types

Adkins series soil and LRSs LHS-1E, LMS-1E, JSC-1A, OPRH4W30, and NUW-LHT-5M were mixed with vermicompost (5% v/v). Mini-tubers were planted once sprouts were approximately 2-cm long. Five tubers were planted in each soil-type (one tuber per pot) in 8.89 x 7.62 cm square pots lined with landscaping fabric and arranged in a randomized format. Plants were scanned with a Phenospex PlantEye600 scanner every 12 hours for the initial 35 days of growth (5 May through 9 June) until plants outgrew the spacing required for scanning. Leaf samples were collected from the upper canopy at 39 DAP and flash frozen for RNA-seq and metabolite analysis. We did not collect leaves from plants grown in NUW-LHT-5M because plants were stunted and did not have leaves. Plants were harvested at 59 DAP and tubers were weighed, flash frozen in liquid nitrogen, lyophilized, and ground to powder with mortar and pestle for later metabolite analysis.

### Soil and tissue analyses

Electroconductivity (EC), pH, moisture, microbial respiration, organic matter content and mineral content, including heavy metals, of soil samples and mineral content of leaf and tuber samples were measured by the Oregon State University Soil Health Lab. For Experiment 1, soil and freeze-dried tissue samples of the same treatment were pooled by measuring and combining equivalent portions of three biological replicates. For experiment 2, four biological replicates were analyzed independently. Additionally, a sample was reserved from the pre-growth batch of each LRS for analysis. EC and pH of soil were determined by suspending the substrate in deionized water in a 1:1 ratio and measured using a Hanna HI5522 instrument (Hanna Instruments, Woonsocket, RI, USA) with either a temperature and pH probe, or an EC probe as appropriate. Microbial respiration was determined by carbon dioxide burst test. Wet substrate was incubated in a vessel with an airtight seal at 23°C for up to 96 h. Carbon dioxide was measured using a Picarro isotopic CO_2_ analyzer (Picarro, Inc., Santa Clara, CA, USA) at 0 h, 24 h, and 96 h. For Experiment 1, minerals were analyzed as follows. Five hundred mg of substrate or 250 mg of plant tissue were acidified using 10 mL nitric acid, then analyzed via microwave digestion using an Anton Paar microwave (Ashland, VA, USA), followed by elemental determination via Inductively Coupled Plasma-Optical Emission Spectrometry (ICP-OES) using an Agilent 5110 instrument (Agilent Technologies, Inc., Santa Clara, CA, USA). For Experiment 2, soil minerals were extracted by adding 20 mL of Mehlich-3 extractant (20 g L^-1^ NH_4_NO_3_, 5.556 mg mL^-1^ NH_4_F, 0.372 mg mL^-1^ EDTA, 11.5 mL L^-1^ glacial acetic acid, and 0.82 mL L^-1^ HNO_3_) to 2 g of sample, shaking for 5 minutes, filtered, then analyzed via ICP-OES. Plant tissue was analyzed using the same method as experiment 1. Note that heavy metal analysis in experiment 1 characterized total elemental content and did not account for bioavailable versus non-bioavailable forms of these metals. Therefore, analysis of these elements in plant tissues will likely provide more meaningful information than those made on soil mixtures. The Mehlich-3 extraction was used in experiment 2 because this method focuses on bioavailable forms of these metals and nutrients.

### Untargeted metabolite analysis using gas chromatography-mass spectrometry (GC-MS)

Tissue extracts were analyzed by GC-MS as previously described [12]. Briefly, 50 mg of dried tissue was extracted using a water:methanol:chloroform (1:2.5:1) solution with ribitol (40 µg mL^-1^) as an internal standard. Extracts were centrifuged to remove cellular debris, then phase separated to isolate aqueous compounds. Fifty µL aliquots were frozen and lyophilized then stored at −80 °C. Samples were derivatized by resuspension of lyophilized material into 20 µL methoxamine HCl (30 mg mL^-1^) dissolved in pyridine and incubating the sample at 37 °C for 90 min with shaking. Then, 40 µL of N-methyl-N-(trimethylsilyl) trifluoroacetamide with 1% chlorotrimethylsilane was added to each sample followed by a 30-min incubation at 37 °C with shaking. One µL of each sample was injected with a 4:1 split for tuber samples and a 10:1 split for leaf samples on an Agilent 7890B GC equipped with an Agilent 5977B MSD (Agilent Technologies). The sample injection order was randomized, and the reagent blank was injected after every three samples to control for potential carryover. Resulting spectra were analyzed using AMDIS [13] and the Agilent Fiehn 2013 GC/MS Metabolomics RTL Library [14] to identify components.

### Targeted metabolite analysis using High-Performance Liquid Chromatography (HPLC)

For HPLC analysis, 100 mg of ground, freeze-dried tuber tissue was extracted in 1 mL of a 50% methanol, 2.5% metaphosphoric acid, 1 mM EDTA extraction solution. Samples were shaken for 10 minutes at room temperature in the dark, then centrifuged at 14,000 x g for 10 minutes. Supernatant was moved to a new tube and re-centrifuged to remove debris. The supernatant was transferred to amber HPLC vials for injection. Metabolites were analyzed using standardized methods as previously published [15]. Standards were prepared in two combination sets: phenolic compounds (ascorbic acid, chlorogenic acid, tyrosine, tryptophan, and phenylalanine) and glycoalkaloids (solanine and chaconine). Phenolic compound standards were prepared with a seven-step 2x serial dilution ranging from 400 to 6.125 mg L^-1^. Serial dilutions of glycoalkaloid standards were prepared at 400, 300, 200, 100, 50, 25, 12.5 and 6.125 mg L^-1^. Each standard was injected in triplicate to generate standard curves used to calculate the compound concentrations in each sample. Compounds were analyzed using an UltiMate 3000 UHPLC system (Thermo Fisher Scientific, Waltham, MA, USA) with an LPG-3400SD quaternary analytical pump, solvent degasser, WPS-3000TSL autosampler, and TCC-3000 column compartment. Compounds were separated using an Acclaim Polar Advantage II C18 column (4.6 x 150 mm, 5 µm) (Dionex, Sunnyvale, CA, USA) using previously published methods [15,16]. Compounds were detected by UV absorbance using a diode array detector DAD 3000 (Thermo Fisher Scientific).

### RNA sequencing

Total RNAs were extracted using a PureLink RNA mini kit (Invitrogen, Waltham, MA, USA), followed by treatment with DNase I (DNA-*free* DNA removal kit (Invitrogen)) to remove traces of genomic DNA. RNA integrity was verified on agarose gel and by analysis on an Agilent Bioanalyzer 2100 (Agilent Technologies, Inc). RNAs were then sent to Novogene for sequencing on an Illumina Novaseq X Plus Series (150-bp, paired-ends) as previously described [17]. Raw .fastq files were run through cutadapt to remove adapter sequences. The DM v6.1 doubled monoploid potato genome and Gene Ontology (GO) term annotations were downloaded from spuddb.uga.edu [18]. Reads were aligned to the genome using HISAT2 [19], sorted using SAMtools [20], and mapped using STRINGTIE [21]. Gene counts were then analyzed in R using the DESeq2 package to generate lists of differentially expressed genes (DEGs), principal component analysis (PCA) plots, and heatmaps [22]. Replicate A of the OPRH4W30 treatment was removed as an outlier based on PCA clustering. Gene set enrichment analysis (GSEA) was performed using the ClusterProfiler package in R [23,24] and used to generate dotplots.

### Statistical analysis

Statistical analyses of GC-MS data were performed using MetaboAnalyst 6.0 (https://www.metaboanalyst.ca/) using default processing settings and data filtering and using the reference feature ribitol for normalization. Features with missing values across replicates were excluded, and a 5% low-variance filter was applied based on the interquantile range. Principle component analyses were generated based on these settings. For other data including harvest metrics, LCMS measurements, PhenospeX data, and PhotsynQ measurements, data was analyzed via ANOVA in R using the ‘dplyr’ and ‘multcompview’ packages. Figures were generated using the ‘ggplot2’, ‘RColorBrewer’ and ‘ggpubr’ packages.

## Results

### Experiment 1: Determination of suboptimal LRS/compost ratio

In this experiment, we tested potato growth on lunar mare simulant LMS-1E amended with varying amounts of vermicompost. Our goal was to find a ratio LRS/compost that supports sufficient plant growth for analysis of growth, physiology, and metabolism but is also suboptimal to investigate the stress response to LRS. In early stages of development, no statistically significant differences in growth and physiology phenotypes recorded by the PlantEye600 scanner were observed (Table S1 & S2). This was likely due to plants emerging from the soil asynchronously, creating large variations within treatment groups at early time points. However, plant growth had leveled out by harvest, and statistically significant differences were observed for multiple growth-related phenotypes (Figure 1). Plants grown in 100% regolith simulant were severely impacted, with reduced total, shoot, root and tuber biomass, as well as harvest index, compared to control plants grown in Adkins soil (Figure 1 & Figure **2**, Table S5). Most notably, the roots of three out of four plants grown in 100% regolith were too small to be weighed on the available balance (less than 0.01 g). Plants grown in 70% regolith simulant were statistically larger than those in 100% regolith in all mass metrics. For some phenotypes,70% LMS grown plants were similar to plants grown in Adkins soil, and even outperformed Adkins soils in tuber mass (Figure 1F). The impact on total tuber mass was also notable, as this translates to the edible biomass of the plant. Plants grown in 85% regolith had a biomass between that of plants grown in 100% regolith simulant and Adkins soil, with no statistically significant difference to either treatment. The variable ratios of compost amendment in the Adkins series control soil had no significant effect on any plant phenotypes. While some photosynthesis related phenotypes measured using PhotosynQ were found to have statistically significant differences, clear patterns across related parameters were not observed (Table S3 & S4).

**Figure 1.**
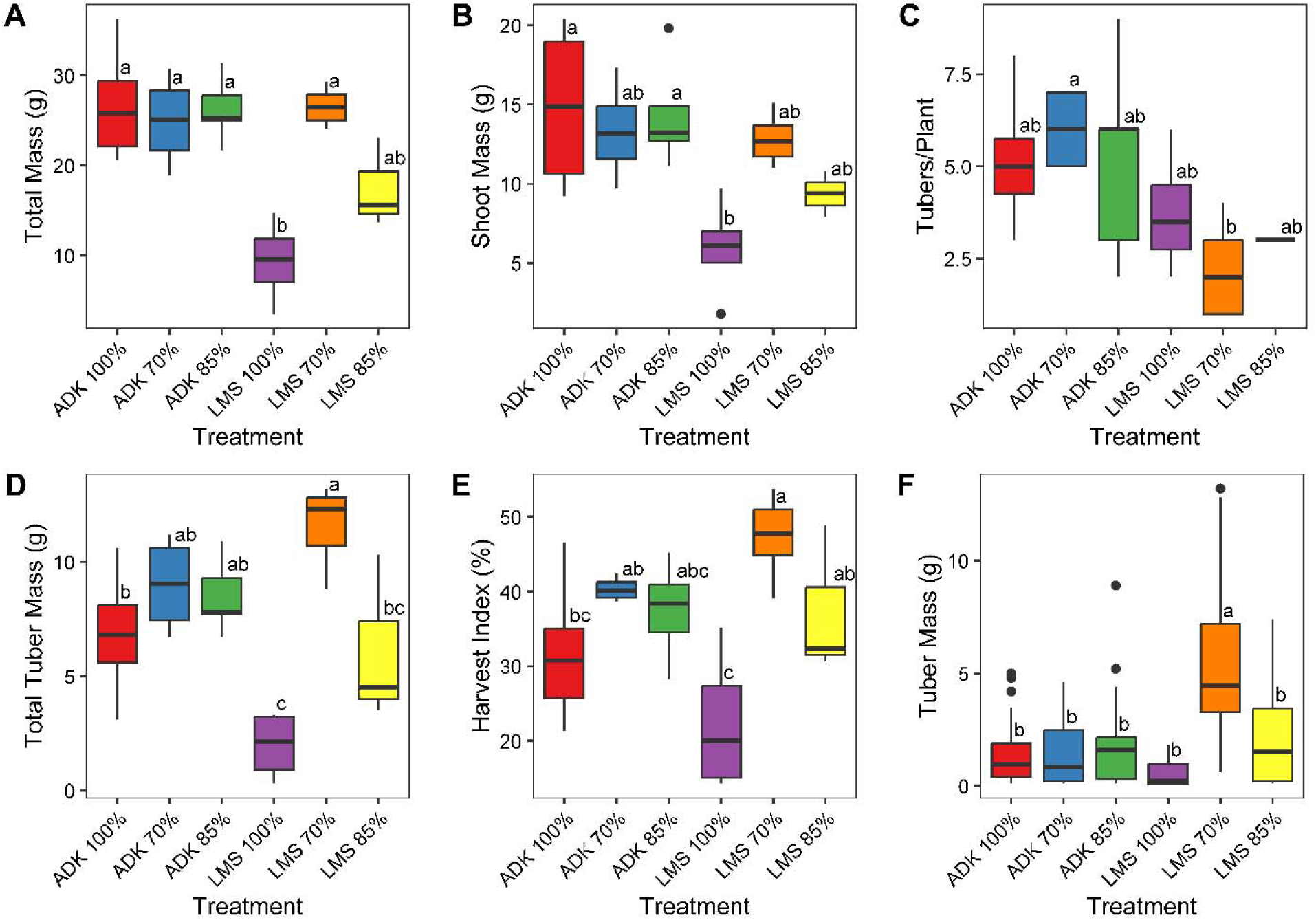
Harvest phenotypes of potato plant variety Modoc grown in variable ratios of LMS-1E and compost. A) Total biomass, B) Shoot biomass, C) Tuber number, D) Total tuber mass, E) Harvest index, and F) Average tuber mass. Shared letters indicate statistically similar groupings as determined by one-way ANOVA and Tukey’s tests.

**Figure 2.**
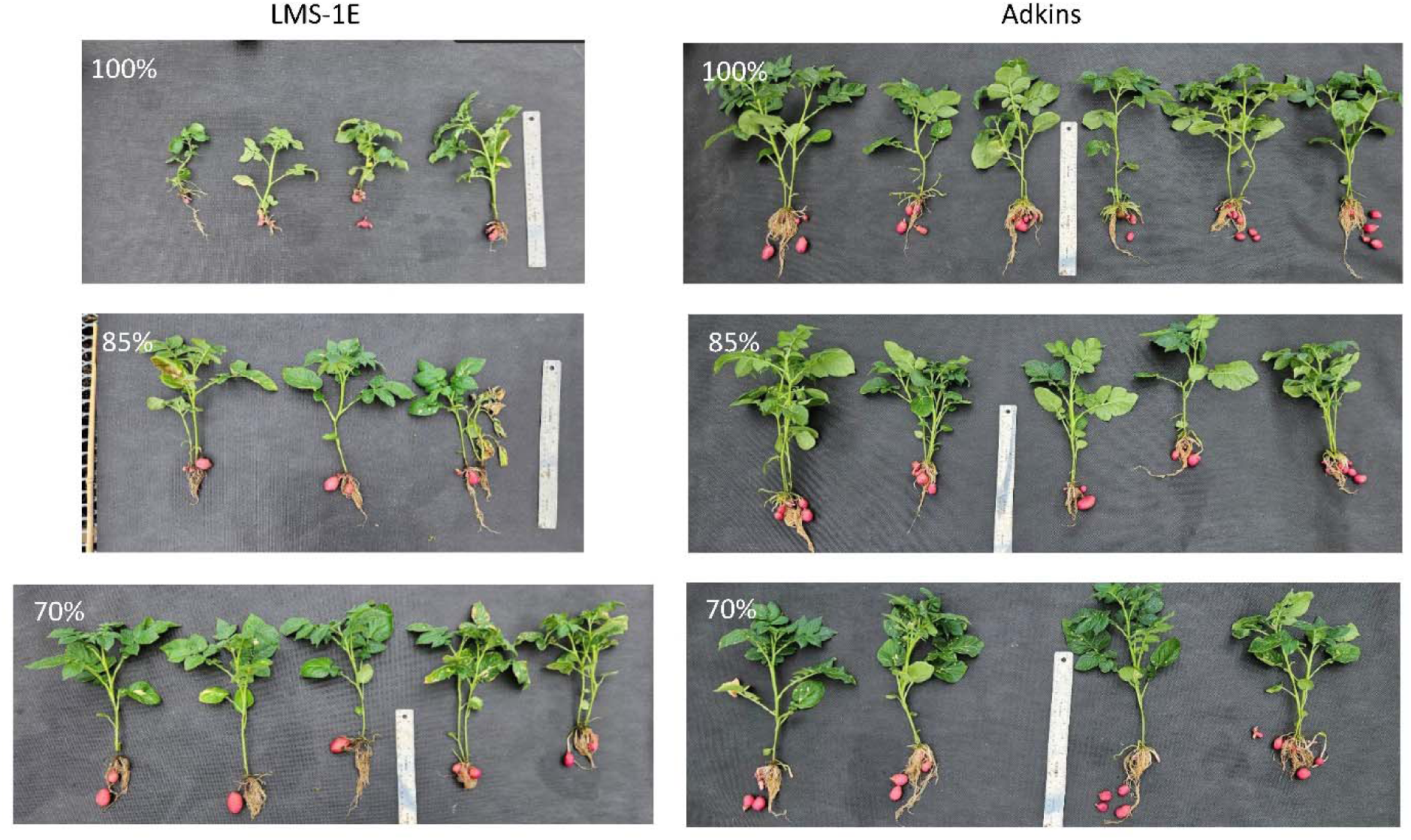
Post-harvest images of potato plants grown in LMS-1E (left) and Adkins soil (right), in 100%, 85%, or 70% regolith to compost volume ratios.

Next, we analyzed soil samples for various metrics to find clues that could explain the phenotypic differences observed. The results presented hereafter are based on analysis of pooled biological replicates, so statistical analysis was not possible. However, several trends are worth noting. Analysis of pre-growth LMS-1E substrate revealed differences in soil characteristics with compost amendment (Table S6). The pH of 100% LMS-1E was 7.9. However, this value decreased as vermicompost content increased, which is expected as pH of vermicompost is between 6 and 7. Increased compost ratio also correlated with an increase in electroconductivity (EC), microbial respiration, and detectable heavy metals. Note that some of these metals most likely come from the Osmocote fertilizer, *i.e.*, Cu, Ni, Zn, as these elements are not present in LMS-1E [4]. Post-growth analysis showed a consistent decrease in pH and increase in EC as compared to pre-growth analyses (Table S6). A drop in pH is expected as plants take up essential ions like K^+^ and Ca^2++^ and compensate by releasing protons into the soil. Post-growth concentrations of heavy metals Al, Cr, Cu, Fe, Ni and Zn were higher than pre-growth. In addition, post-growth microbial respiration decreased compared to pre-growth analysis for 85% and 70% regolith simulants, while 100% regolith had slightly increased levels. It should be noted that microbial respiration differed between 85% and 70% regolith treatment, despite containing the same amount of non-sterilized compost. This is likely due to the method of assessment, which involves incubating the soil for 24-96 h, during which the microbes reproduce at different rates based on the quantity of compost in the substrate.

To further understand what factors might have led to underdeveloped roots and overall decreased plant growth in 100% regolith simulant, we quantified heavy metal contents in leaves and tubers (Table S7). The most striking observation was the large accumulation of copper in leaves compared to all other treatments, and to a lesser extent in tubers, of plants grown in 100% regolith simulant compared to plants grown in Adkins soil and 70% and 85% regolith simulant, suggesting that copper concentrations available to plants were at toxic levels compared to Adkins.

### Experiment 2: Effect of various lunar regolith simulant types

#### The growth of potato plants varies depending on LRS type

In Experiment 1, we found a relatively linear relationship between added compost and improvement of plant growth. This information was used to inform the design of the second experiment. Stunting caused by 100% regolith simulant limits analyses due to lack of sample, while 70% and 85% simulant may mask stress responses caused by the regolith. As such, we used a 95:5 v/v ratio of regolith to compost in this second experiment. In addition, we planted actively sprouting mini-tubers (approximately 2-cm long sprouts) to decrease variability in plant development due to asynchronous emergence of shoots from tubers. We selected three lunar highland simulants (LHS-1E, OPRH4W30, and NUW-LHT-5M) and two lunar mare simulants (JSC-1A and LMS-1E). Plants grown in lunar highland simulant NUW-LHT-5M were the most severely affected, with statistically significant reduction in plant size and tuber biomass (Figures 3 & 4). This can be partly explained by the severely delayed emergence of plants observed in this LRS (approximately 24 DAP compared to 4-8 DAP for other types of LRS). Additionally, only three out of five plants grown on NUW-LHT-5M grew large enough to be detected by the Phenospex scanner and produced tubers (Tables S8 & S9). Statistically significant reduction in tuber mass was also observed in all other LRS types compared to Adkins soil (Figure 4), while plant size was statistically reduced only in lunar highland simulant OPRH4W30 compared to Adkins when analyzing digital canopy biomass (Figure 3). Colorimetric indices did not have any statistical differences between treatments (Table S8).

**Figure 3.**
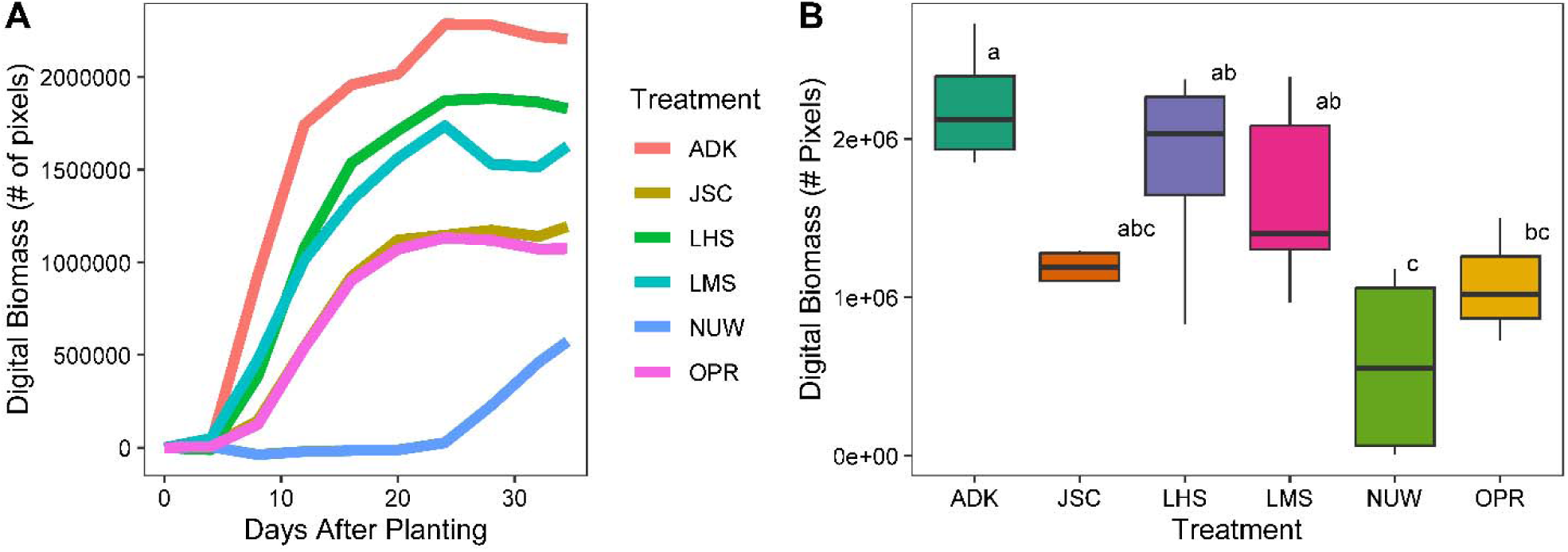
(A) Digital above-ground biomass across the first 34 days after planting (DAP) of potato plants grown in Adkins soil (ADK) and five regolith simulants; JSC-1A (JSC), LHS-1E (LHS), LMS-1E (LMS), NUW-LHT-5M (NUW), and OPRH4W30 (OPR). Data was collected via the Phenospex PlantEye600 scanner. (B) Boxplot of final above-ground digital biomass at 34 DAP. All substrates were amended with 5% compost by volume. Shared letters indicate statistically significant groupings determined by ANOVA followed by Tukey’s post-hoc. N=5 except NUW-LHT-5M where N=3.

**Figure 4.**
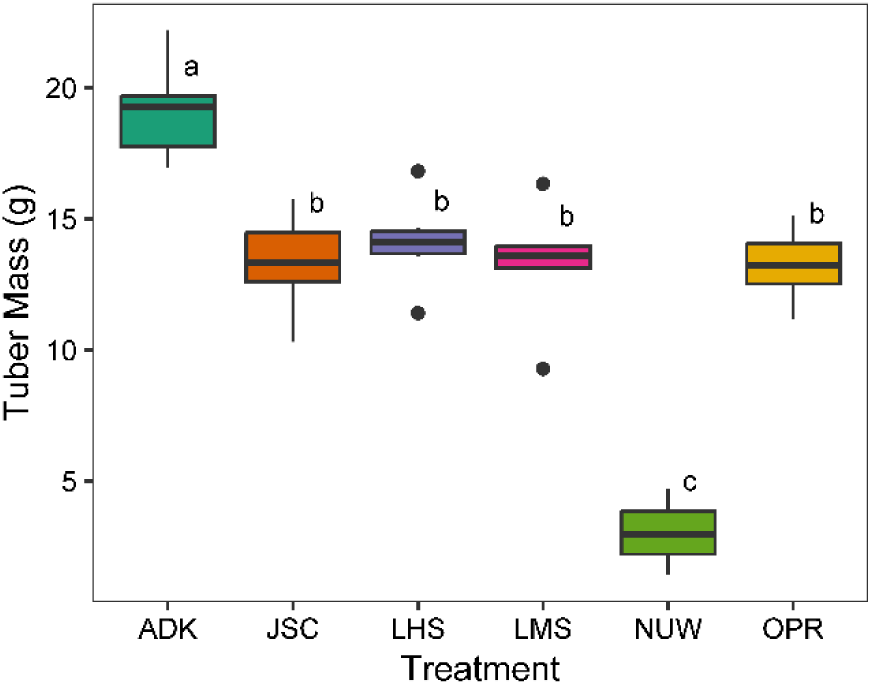
Harvested tuber mass of plants. Tubers were harvested at 59 DAP from plants grown in Adkins soil (ADK) and five regolith simulants; JSC-1A (JSC), LHS-1E (LHS), LMS-1E (LMS), NUW-LHT-5M (NUW), and OPRH4W30 (OPR). Shared letters a, b and c indicate statistically significant groupings determined by ANOVA followed by Tukey’s post-hoc. N=5 except NUW-LHT-5M where N=3.

#### The potato transcriptome response differs between LRSs

To better understand the molecular changes that led to the observed phenotypes, we analyzed the transcriptome of leaves harvested from plants grown on lunar mare simulants JSC1-A and LMS-1E, lunar highland simulants OPRH4W30 and LHS1-E, and the Adkins terrestrial soil. A total of 972,140,076 raw reads were obtained (Table S10). Differential gene expression analysis between Adkins and each LRS revealed varying numbers of differentially expressed genes (DEGs) (p-adjusted value < 0.05) (Figure 5, Table S11). Comparison between lunar highland simulant OPRH4W30 and Adkins produced the lowest number of DEGs at 124, while 958, 1397, and 360 genes were differentially expressed when comparing lunar mare simulants JSC1-A and LMS1-E and lunar highland simulant LHS1-E, respectively, with Adkins. The majority of DEGs in response to lunar mare simulants JSC-1A and LMS-1E were unique to each of these LRSs, whereas most DEGs in response to lunar highland simulant LHS-1E were common with those found in response to lunar mare simulant LMS-1E. Gene set enrichment (GSE) analysis of lunar mare simulants JSC1-A and LMS-1E and lunar highland simulants LHS1-E and OPRH4W30 revealed enrichment of 6, 20, 6, and 7 gene ontology terms, respectively (p-adj < 0.05) (Figure 6, Table S12). Many affected genes are related to various stress response and signaling pathways. Notably, all four LRSs affected genes related to ‘kinase activity’ (GO:0016301). Four pathways were enriched in LMS-1E, LHS-1E, and OPRH4W30: ‘extracellular region’ (GO:0005576), ‘protein modification process’ (GO:0036211), ‘protein metabolic process’ (GO:0019538), and ‘enzyme regulator activity’ (GO:0030234). The pathways ‘cell wall’ (GO:0005618) was enriched in both LMS-1E and OPRH4W30. LHS-1E, JSC-1A, and OPRH4W30 shared one enriched process: ‘signal transduction’ (GO:0007165). Thirteen pathways were uniquely enriched by LMS-1E: ‘DNA-binding transcription factor activity’ (GO:0003700), ‘DNA binding’ (GO:0003677), ‘cytosol’ (GO:0005829), ‘nucleobase-containing compound metabolic process’ (GO:0006139), ‘response to chemical’ (GO:0042221), ‘catalytic activity’ (GO:0003824), ‘endoplasmic reticulum’ (GO:0005783), ‘nuclease activity’ (GO:0004518), ‘signaling receptor activity’ (GO:0038023), ‘thylakoid’ (GO:0009579), ‘RNA binding’ (GO:0003723), ‘hydrolase activity’ (GO:0016787) and ‘membrane’ (GO:0016020). Three pathways were uniquely enriched by JSC-1A: ‘carbohydrate binding’ (GO:0030246), ‘molecular function’ (GO:0003674), and ‘response to external stimulus’ (GO:0009605). LHS-1E and OPRH4W30 did not uniquely enrich any pathway.

**Figure 5.**
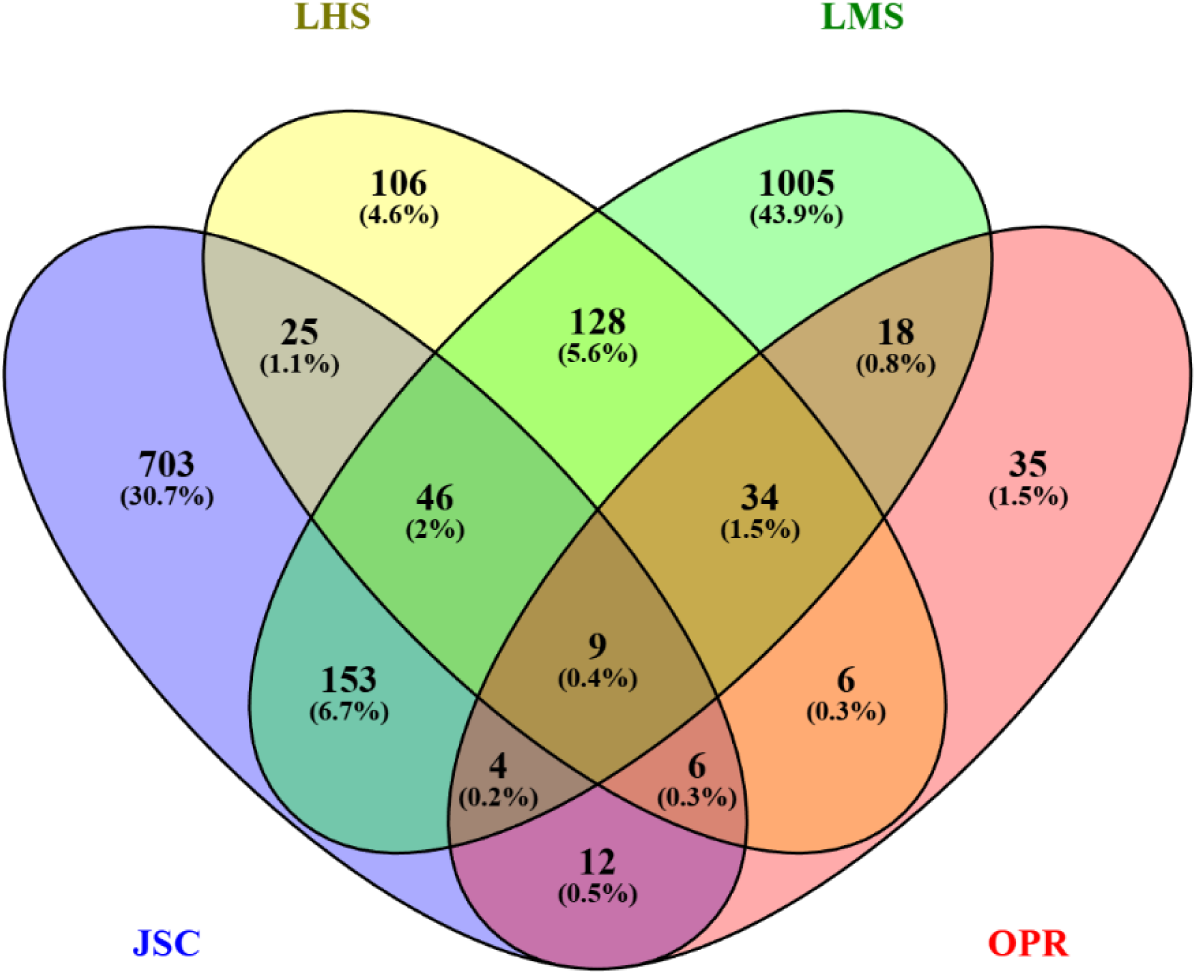
Venn diagram of differentially expressed genes (p_adj_ < 0.05) of plants grown in LHS- 1E (LHS), LMS-1E (LMS), JSC-1A (JSC), and OPRH4W30 (OPR) compared with the terrestrial soil Adkins.

**Figure 6.**
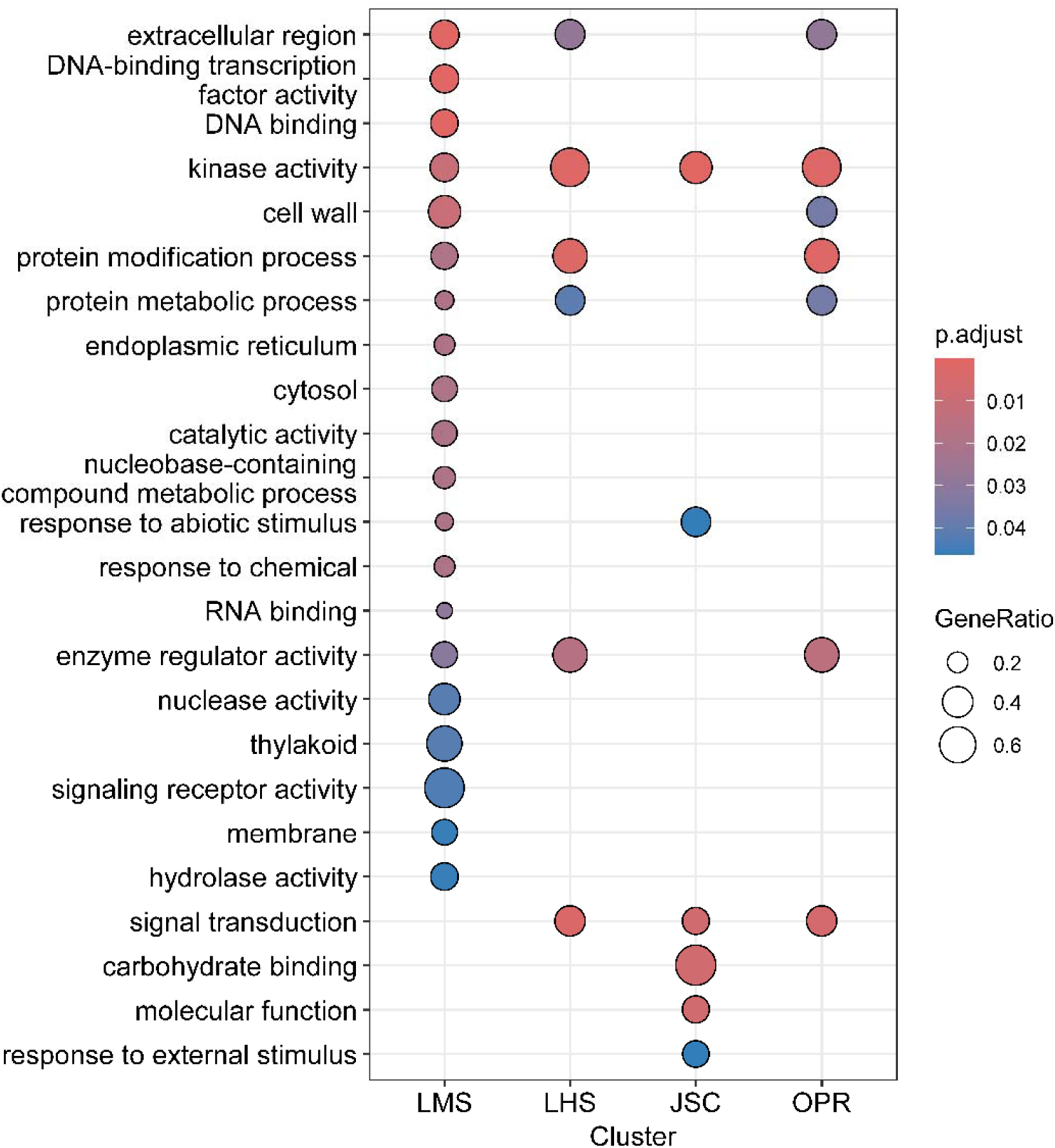
Gene Set Enrichment (GSE) analysis of differentially expressed genes between Adkins soil treatment and LRS. LMS-1E (LMS), LHS-1E (LHS), JSC-1A (JSC), OPRH4W30 (OPR). Dotplots display the gene ratio (count of enriched genes/count of total genes in pathway) of various GO terms based on comparison to Adkins control soil.

#### Metabolites and minerals profiles are unique to each LRS

We complemented transcriptome analysis with metabolite analysis in both leaf and tuber tissues. Analysis of leaf metabolites by GC-MS revealed unique metabolic profiles of plants grown in LRS as compared to Adkins soil (Figures 7, Table S13). Compounds contributing to these differences include sucrose and fructose as well as other nutrients such as citric acid (Figure 8). Analysis of tuber metabolites by GC-MS revealed only minor changes between treatments (Figure 8, Table S14). In addition, analysis of phenolic compounds and two major glycoalkaloids via HPLC-UV/VIS did not reveal significant changes in 6 out of 7 compounds analyzed (Figure 9). Tyrosine concentrations increased significantly in plants grown in lunar mare simulant JSC-1A compared to Adkins, but did not increase significantly in lunar mare simulant LMS-1E and lunar highland simulants LHS-1E and OPRH4W30. Heavy metal analysis of tubers revealed increased accumulation of Cu and Zn in all LRS grown-tubers, though not statistically significant in some cases. Increased accumulation of Al in tubers grown in lunar highland simulants LHS-1E and OPRH4W30 was also observed (Table 2, Table S15).

**Figure 7.**
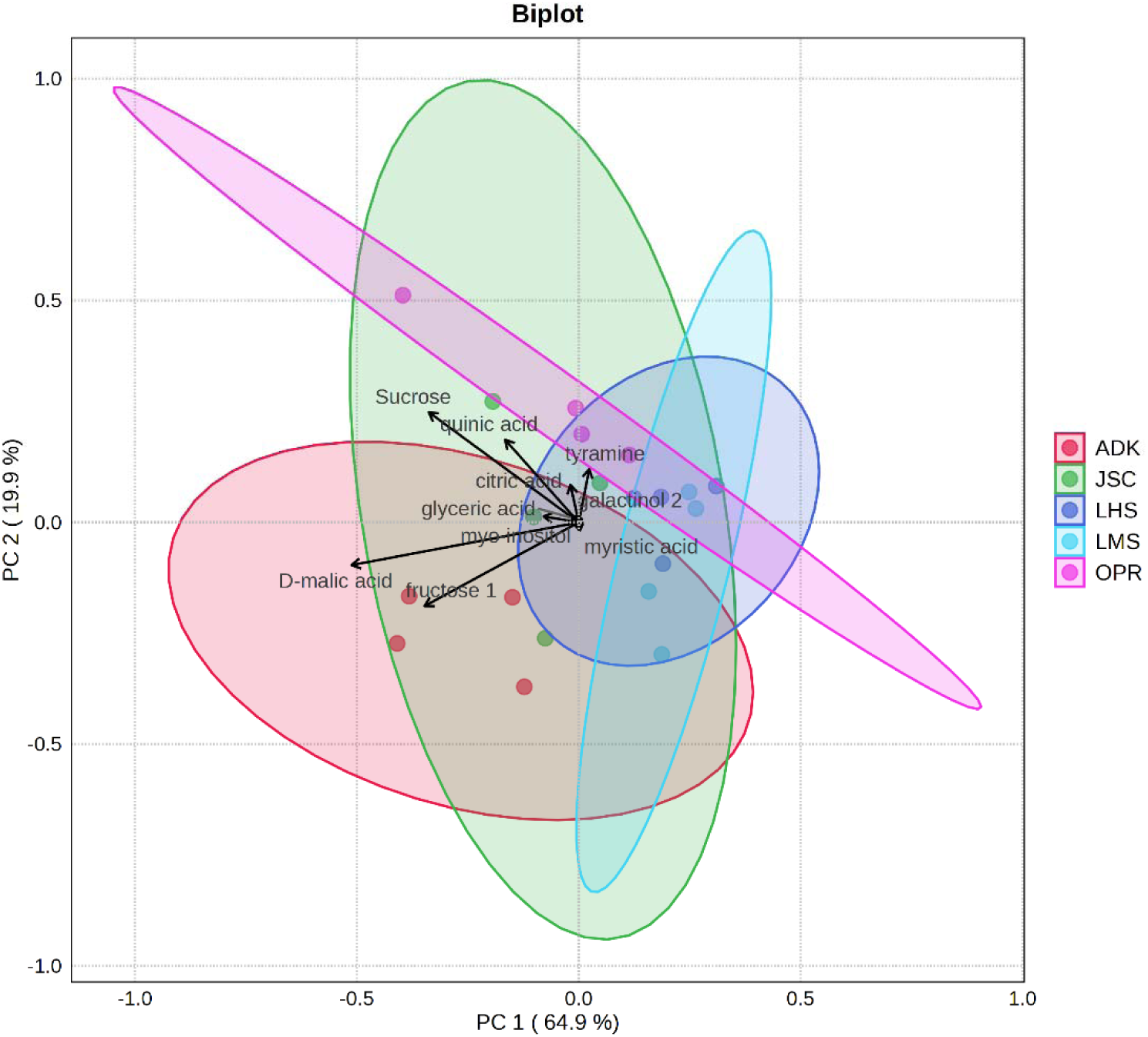
PCA biplot of GC-MS metabolite analysis of leaves collected at 39 DAP from potato plants grown in Adkins soil (ADK) and four lunar regolith simulants. JSC-1A (JSC), LHS-1E (LHS), LMS-1E (LMS), and OPRH4W30 (OPR).

**Figure 8.**
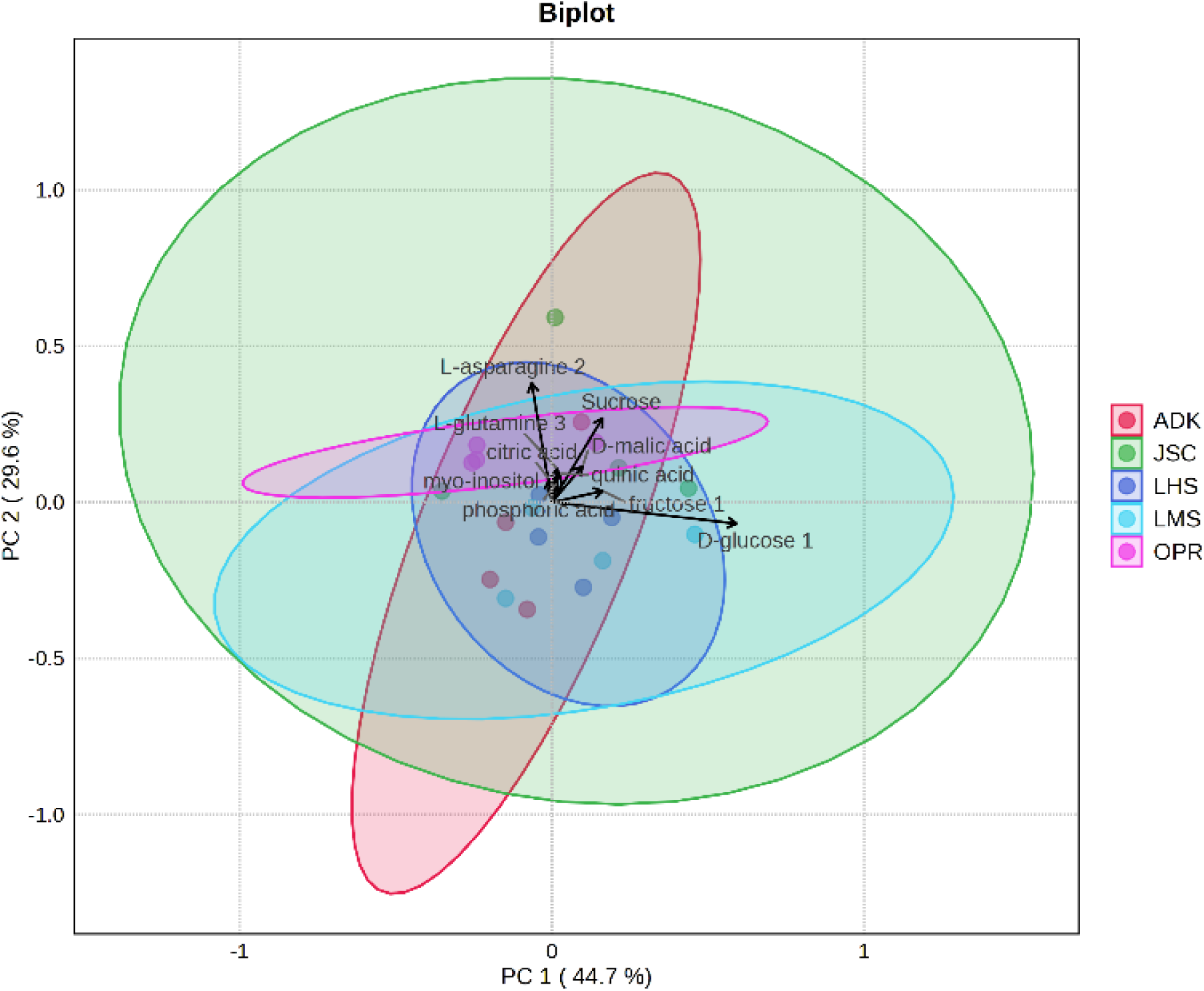
PCA biplot of GC-MS metabolite analysis of tubers harvested at 59 DAP from potato plants grown in Adkins soil (ADK) and four lunar regolith simulants; JSC-1A (JSC), LHS-1E (LHS), LMS-1E (LMS), and OPRH4W30 (OPR).

**Figure 9.**
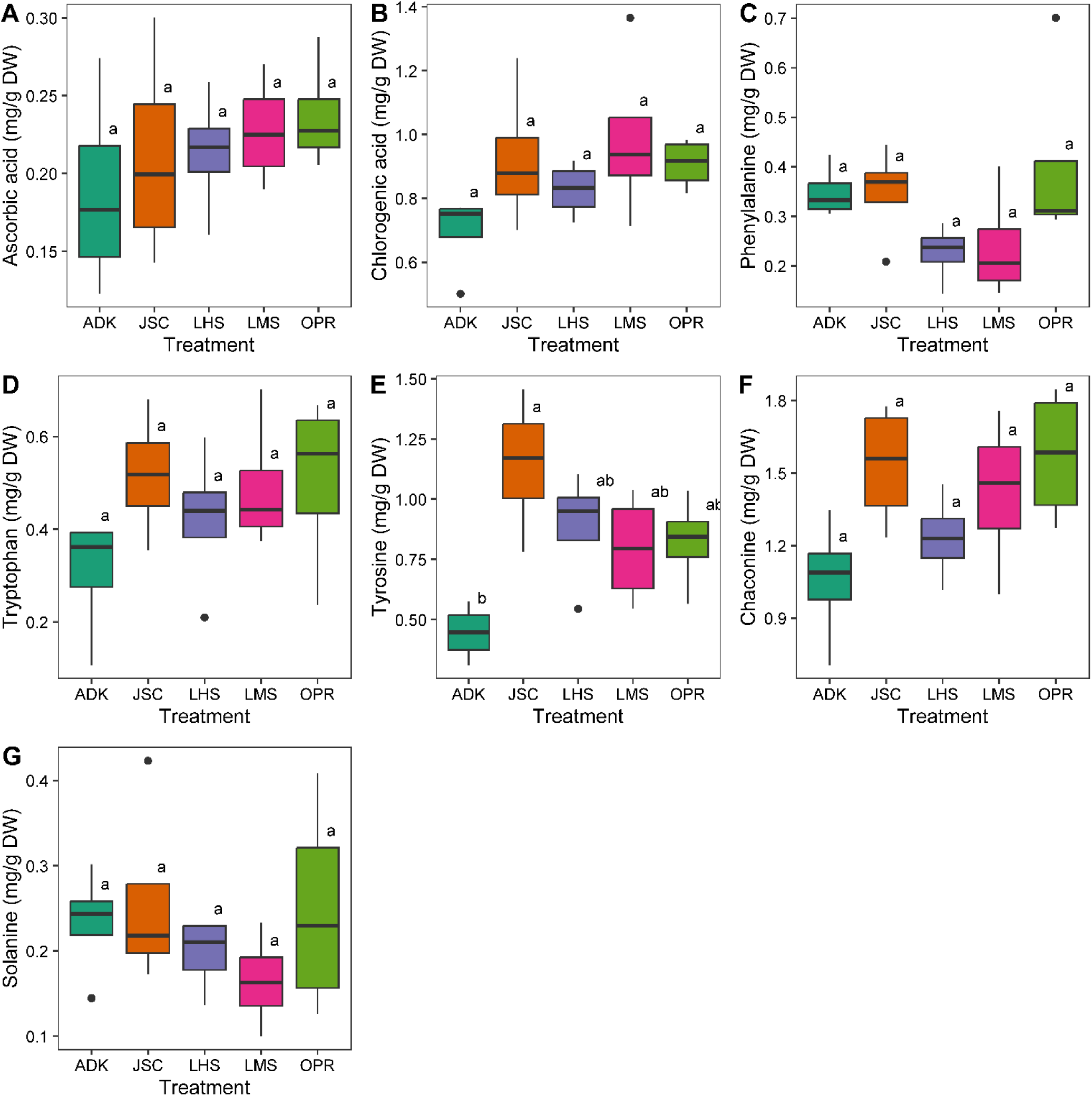
Quantities of targeted compounds extracted from harvested tubers of potato plants grown in Adkins soil (ADK) and five lunar regolith simulants. JSC-1A (JSC), LHS-1E (LHS), LMS-1E (LMS), and OPRH4W30 (OPR). (A) ascorbic acid, (B) chlorogenic acid, (C) phenylalanine, (D) tryptophan, and (E) tyrosine, (F) chaconine, and (G) solanine. Shared letters indicate statistically significant groupings determined by ANOVA followed by Tukey’s post-hoc.

#### Analysis of soils and tuber heavy metal content

Soil analysis revealed differences between LRSs and Adkins soil (Table 1 or Table S16). All LRSs had higher moisture content, indicating reduced drainage capability. Lunar mare simulants LMS1-E and JSC-1A and lunar highland simulant, LHS1-E all experienced a reduction in pH from 7.5-7.72 range to ∼6.5, whereas lunar highland simulant OPRH4W30 remained ∼7.5, compared to pre-growth samples (Table S17). Electroconductivity decreased in lunar mare simulants LMS1-E and JSC-1A and increased in lunar highland simulants LHS-1E and OPRH4W30 compared to pre-growth samples. Elemental availability was also altered for some elements, such as Ca, K, Mg, and Zn, post-growth. However, this is consistent with changes in pH, which determines the solubility of many nutrients.

**Table 1.**
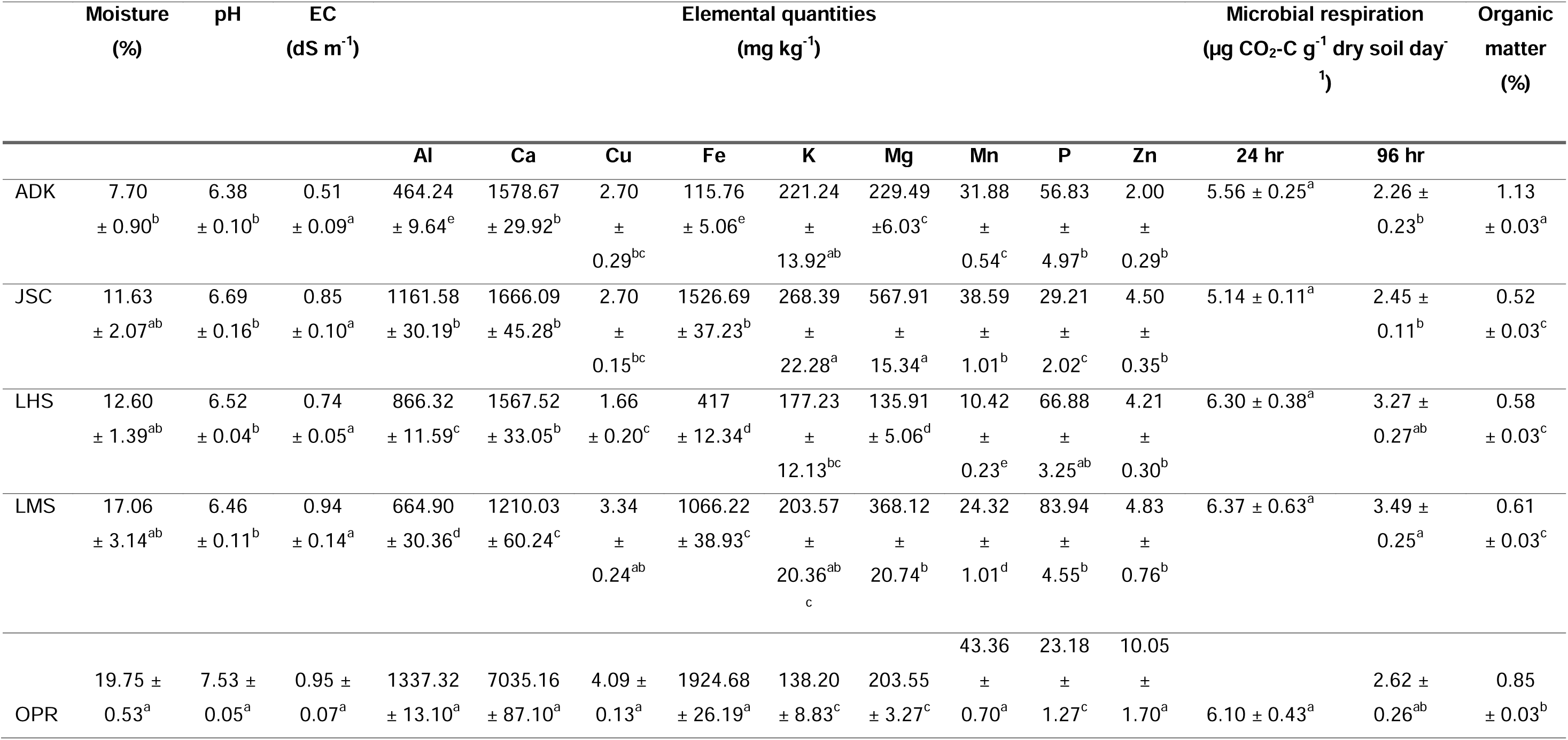
Post-growth soil characteristics of soil substrates amended with 5% v/v vermicompost. Data are means ± SE (N =4). Superscript letters indicate statistically similar groupings according to ANOVA followed by Tukey’s post-hoc. Analysis was performed in R.

**Table 2.** Heavy metal content of tubers grown in various LRS with 5% v/v vermicompost, determined by ICP-OES. Letters indicate statistically similar groupings determined by one-way ANOVA and Tukey’s post-hoc, performed in R. N=4. As, Cd, Co, Cr, Ni, Pb were also analyzed but found below quantifiable levels in all samples.

| Soil type | Heavy metal content<br>(mg kg<sup>-1</sup> dry weight) |  |  |
| --- | --- | --- | --- |
|  | Al | Cu | Zn |
| ADK | 55.9 ± 3.03<sup>b</sup> | 2.8 ± 0.32<sup>b</sup> | 6.6 ± 0.31<sup>b</sup> |
| JSC | 59.3 ± 6.42<sup>b</sup> | 6.0 ± 0.60<sup>a</sup> | 12.3 ± 1.08<sup>a</sup> |
| LHS | 144.6 ± 11.60<sup>a</sup> | 4.7 ± 1.08<sup>ab</sup> | 9.5 ± 0.63<sup>ab</sup> |
| LMS | 50.8 ± 3.74<sup>b</sup> | 5.1 ± 0.00<sup>ab</sup> | 11.4 ± 0.51<sup>a</sup> |
| OPR | 185.1 ± 26.88<sup>a</sup> | 6.0 ± 0.32<sup>ab</sup> | 12.0 ± 0.64<sup>a</sup> |

## Discussion

The goal of this study was to assess the effects of LRS on potato growth and physiology and to characterize the underlying molecular changes. Because LRSs do not contain any nitrogen, we provided plants with this macronutrient along with other macro and micronutrients via fertilization with the slow-release fertilizer Osmocote. Growth was severely affected when potato plants were grown in 100% lunar simulant LMS-1. Symptoms included stunting and underdeveloped roots, which are typical of plants with excess copper [25,26]. Interestingly, leaf and tuber tissues of plants grown in 100% LMS-1E accumulated higher copper levels compared to Adkins control. The only notable source of copper was Osmocote, as copper is not present in LMS-1E [27]. Excess aluminum can also lead to similar phenotypes [28]. However, we did not observe any substantial differences in leaf and tuber Al concentrations between plants grown in 100% LMS-1E and Adkins. This may be due to the low solubility of aluminum at pH measured in the substrates used in this study, which were consistently above 5. Aluminum becomes an issue at pH below 5 that increases solubility [29]. The negative effect of 100% LMS-1E was alleviated by adding compost to the growth substrate. If copper toxicity is the reason for the stunting phenotype, the addition of organic matter would provide binding sites for copper [30], thereby alleviating hyperaccumulation. These types of interactions should therefore be considered when designing fertilizer management plans for growing crops in lunar regolith that had little to no biological amendments.

Potato plants grown on different lunar mare and lunar highland regolith simulants amended with 5% compost were all negatively affected in canopy size and tuber yield compared to those grown in Adkins. Therefore, LRS is stressful for potato plants, which responded to this stressful environment by modifying the expression of genes involved in photosynthesis, stress responses, signaling, and metabolism of terpenes and flavonoids. We have several hypotheses to help explain these negative phenotypes. First, we cannot rule out that copper and possibly aluminum toxicity was completely alleviated by adding compost. Indeed, heavy metals analysis in tubers showed that copper concentrations were significantly higher in lunar mare highland JSC-1A compared to Adkins, and aluminum concentrations were higher in lunar highland simulants LHS-1E and OPRH4W30, indicating higher uptake of these heavy metals in LRS than in Adkins. Copper is also moderately increased in all other LRS, though not statistically different according to ANOVA. Unfortunately, we were unable to confirm these results in leaf tissues as plants had senesced at harvest, preventing us from collecting leaf samples. It remains unclear how aluminum could accumulate in tubers grown on lunar highland simulants as pH was above 5. Second, the presence of glassy agglutinates in lunar highland simulants OPRH4W30 and NUW-LHT-5M may be particularly problematic for plant growth [31,32]. Lunar mare simulant JSC-1A, despite lacking in agglutinates, does contain a high glass fraction [33] and caused similar issues as OPRH4W30. These sharp micro-particles may cause abrasions on root/stolon/tuber epidermis. Given the findings of the transcriptomic analysis, these microabrasions are likely being perceived by the plant as herbivory by nematodes and other soil-borne insects (Figure 6). Third, high levels of compaction, cementation, small particle size and higher density of lunar highland simulant NUW-LHT-5M and lunar mare simulant JSC-1A after watering might also have negatively affected plant growth and development. This is consistent with the higher moisture content recorded in LRSs vs the Adkins soil, and is likely a result of high particle density and small particle size. Finally, it is likely that all together these factors caused the stunting observed.

Growth in LRS did not significantly alter potato tuber metabolite profiles, indicating negligible changes to nutritional content. Additionally, stress induced by the LRS did not cause any changes in glycoalkaloid levels. These findings support that tuber nutritional quality is maintained in LRS-grown tubers, despite lower potential yields. Should this limitation prove true with larger-scale growth of mature plants, agricultural modules may need to be larger than currently predicted to produce required quantities of edible matter. Heavy metal analysis revealed mild accumulation of Zn and Cu in tubers grown in LRS. In addition, plants grown in highland-type LRS were found to accumulate aluminum, which is not of immediate concern for human consumption purposes, but could contribute to long-term negative impacts on plant production.

## Conclusions

From these observations, we conclude that LRS are clearly stressful on potato plants, given the reduction in overall and edible biomass, and the changes to stress response related gene expression. However, significant changes were not found in metabolite profiles of leaves or tubers, indicating that nutrition was not impacted by growth in LRS. Finally, addition of vermicompost to LMS-1E improves growth of potato. This highlights that remediation of lunar regolith via the addition of biomatter is possible, which bodes well for later generations of plant growth in lunar regolith as biomatter builds up over the course of the first generations, though the lack of organic matter presents an issue for early stages of bioregenerative life support development.

## Declarations

### Ethics approval and consent to participate

Not applicable

### Consent for publication

Not applicable

### Availability of data and materials

The raw sequence reads have been deposited in the Open Science Data Repository under doi 10.26030/w56j-2d69.

### Authors’ contributions

DH: design of the work, acquisition, analysis, and interpretation of data, wrote first draft of the manuscript; AL: data acquisition; MK: data acquisition; SC: data acquisition; PJ: funding acquisition, design of the work, reviewed and edited the manuscript; JCA: data acquisition, reviewed and edited the manuscript; AG: conception, funding acquisition, design of the work, analysis and interpretation of data, reviewed and edited the manuscript. All authors have approved the submitted manuscript.

## Acknowledgements

We thank Matthew Barrow at CSS Farms for providing minitubers of Modoc.

