## Supplementary Materials for "Growth and molecular responses of potato to lunar regolith simulants"

### Additional Materials

**Additional Table S2**. Average values with standard deviation of metrics recorded by Phenospex PlantEye600 scanner at 32 DAP, 18 days after start of scanning, of plants grown in Adkins soil or LMS-1E, supplemented with vermicompost in 70:30, 85:15, or 100:0 soil to compost ratio. ANOVA performed in R revealed no statistically significant groupings amongst treatments in any metric.

| Treatment | Digital Biomass (mmÂ³) | Leaf Area | Hue | NDVI | NPCI | PSRI | Greenness |
| --- | --- | --- | --- | --- | --- | --- | --- |
| Adkins 100% | 348837.50 ± 209702.26 ^a^ | 3125.76 ± 1616.81 ^a^ | 103.95 ± 6.75 ^a^ | 0.621 ± 0.073 ^a^ | 0.094 ± 0.022 ^a^ | 0.056 ± 0.034 ^a^ | 0.214 ± 0.035 ^a^ |
| Adkins 85% | 343960.68 ± 226103.07 ^a^ | 4081.16 ± 1742.94 ^a^ | 103.87 ± 12.57 ^a^ | 0.561 ± 0.205 ^a^ | 0.053 ± 0.009 ^a^ | 0.024 ± 0.004 ^a^ | 0.180 ± 0.065 ^a^ |
| Adkins 70% | 275665 ± 218456.21 ^a^ | 3536.51 ± 2519.16 ^a^ | 112.22 ± 2.44 ^a^ | 0.700 ± 0.025 ^a^ | 0.064 ± 0.020 ^a^ | 0.023 ± 0.006 ^a^ | 0.242 ± 0.035 ^a^ |
| LMS-1E 100% | 264856.5 ± 139810.04 ^a^ | 3383.30 ± 1147.25 ^a^ | 112.25 ± 2.31 ^a^ | 0.679 ± 0.017 ^a^ | 0.058 ± 0.015 ^a^ | 0.022 ± 0.008 ^a^ | 0.221 ± 0.023 ^a^ |
| LMS-1E 85% | 617795.95 ± 325939.28 ^a^ | 5655.34 ± 2184.15 ^a^ | 107.00 ± 9.87 ^a^ | 0.629 ± 0.112 ^a^ | 0.081 ± 0.037 ^a^ | 0.047 ± 0.046 ^a^ | 0.198 ± 0.053 ^a^ |
| LMS-1E 70% | 642838 ± 386465.63 ^a^ | 5724.90 ± 2816.72 ^a^ | 112.41 ± 1.23 ^a^ | 0.690 ± 0.021 ^a^ | 0.062 ± 0.013 ^a^ | 0.019 ± 0.002 ^a^ | 0.232 ± 0.022 ^a^ |

**Additional Table S4**. Average values with standard deviation of metrics recorded by PhotosynQ scanner at 63 DAP of plants grown in Adkins soil or LMS-1E, supplemented with vermicompost in 70:30, 85:15, or 100:0 soil to compost ratio. Letters indicate statistically similar groupings determined by ANOVA performed in R. Metrics include magnitude of electrochromic shift (ECS_t_), proton conductivity (gH^+^), quantum yield (Φ_II_), linear electron flow (LEF), non-regulatory energy dissipation (Φ_NO_), non-photochemical quenching (Φ_NPQ_ and NPQ_t_), relative chlorophyll via special products analysis division (SPAD), and the redox states of photosystem 1 (PS1).

| Treatment | ECS_t_ | gH^+^ | Φ_II_ | LEF | Φ_NO_ | Φ_NPQ_ | NPQ_t_ | SPAD | PS1 active centers | PS1 open centers | PS1 over reduced centers | PS1 oxidized centers |
| --- | --- | --- | --- | --- | --- | --- | --- | --- | --- | --- | --- | --- |
| Adkins 100% | 0.011 ± 0.005^a^ | 117.188 ± 49.783 ^a^ | 0.590 ± 0.051 ^a^ | 49.281 ± 28.839 ^b^ | 0.232 ± 0.036 ^a^ | 0.179 ± 0.073^b^ | 0.827 ± 0.442^b^ | 44.972 ± 7.984^a^ | 2.621 ± 0.800 ^a^ | 0.416 ± 0.241 ^a^ | 0.428 ± 0.220 ^a^ | 0.156 ± 0.127 ^b^ |
| Adkins 85% | 0.013 ± 0.005^a^ | 93.243 ± 38.438 ^a^ | 0.469 ± 0.134^b^ | 88.350 ± 47.793 ^a^ | 0.219 ± 0.061 ^a^ | 0.313 ± 0.165 ^a^ | 1.640 ± 1.026 ^a^ | 44.691 ± 12.255^a^ | 2.851 ± 1.226 ^a^ | 0.367 ± 0.260 ^a^ | 0.313 ± 0.295 ^a^ | 0.320 ± 0.199 ^a^ |
| Adkins 70% | 0.010 ± 0.00^a^ | 109.127 ± 42.151^a^ | 0.505 ± 0.167 ^ab^ | 65.395 ± 33.839 ^ab^ | 0.204 ± 0.057 ^a^ | 0.215 ± 0.114 ^ab^ | 1.046 ± 0.677 ^ab^ | 44.523 ± 12.357^a^ | 2.588 ± 0.798 ^a^ | 0.520 ± 0.243 ^a^ | 0.204 ± 0.154 ^a^ | 0.238 ± 0.151 ^ab^ |
| LMS-1E 100% | 0.007 ± 0.00^a^ | 42.604 ± 349.92 ^a^ | 0.611 ± 0.040 ^ab^ | 45.486 ± 35.354 ^b^ | 0.239 ± 0.035 ^a^ | 0.151 ± 0.063^b^ | 0.676 ± 0.358 ^ab^ | 41.956 ± 13.173^b^ | 2.569 ± 0.683 ^a^ | 0.462 ± 0.234 ^a^ | 0.419 ± 0.275 ^a^ | 0.119 ± 0.089 ^ab^ |
| LMS-1E 85% | 0.012 ± 0.007 ^a^ | 101.000 ± 56.668 ^a^ | 0.510 ± 0.170 ^a^ | 60.030 ± 34.592 ^ab^ | 0.194 ± 0.062 ^a^ | 0.216 ± 0.127 ^ab^ | 1.143 ± 0.857 ^ab^ | 43.967 ± 13.662^a^ | 2.575 ± 0.749 ^a^ | 0.497 ± 0.301 ^a^ | 0.267 ± 0.197 ^a^ | 0.193 ± 0.155 ^ab^ |
| LMS-1E 70% | 0.009 ± 0.003 ^a^ | 130.171 ± 39.428 ^a^ | 0.605 ± 0.046 ^a^ | 39.358 ± 16.257 ^b^ | 0.250 ± 0.025 ^a^ | 0.145 ± 0.050^b^ | 0.593 ± 0.235^b^ | 47.436 ± 2.746^a^ | 2.909 ± 0.938 ^a^ | 0.519 ± 0.237 ^a^ | 0.387 ± 0.228 ^a^ | 0.095 ± 0.063 ^b^ |

**Additional Table S6.** Soil analysis results of regolith simulant LMS-1E mixed with variable ratios of compost, before and after growth of Modoc. Heavy metals As, Cd, Co, and Pb were also analyzed but were below quantifiable levels. Pre-growth data were obtained from one batch of soil sample per treatment. Post-growth data were based on the analysis of pooled biological replicated samples for each treatment.

| Soil Type | pH | EC  (dS m^-1^) | Microbial respiration  (µg CO_2_-C g^-1^ dry soil day^-1^) | | Heavy Metal  (mg kg^-1^) | | | | | |
| --- | --- | --- | --- | --- | --- | --- | --- | --- | --- | --- |
| *Pre-Growth* |  |  | 24 hr | 96 hr | Al | Cr | Cu | Fe | Ni | Zn |
| 100% LMS | 7.9 | 0.73 | 2 | 1 | 3077 | 14.9 | 24.5 | 10327 | 43.6 | 18.1 |
| 85% LMS | 7.37 | 1.24 | 16 | 8 | 3269 | 16.7 | 25.6 | 10989 | 46.1 | 20 |
| 70% LMS | 7.06 | 2.03 | 25 | 14 | 3267 | 18.6 | 30.1 | 11870 | 49.9 | 22.8 |
| *Post-Growth* |  |  |  |  |  |  |  |  |  |  |
| 100% LMS | 6.39 | 0.92 | 4 | 2 | 3609 | 20 | 28 | 12582 | 54 | 35 |
| 85% LMS | 5.22 | 2.91 | 6 | 5 | 3471 | 18 | 29 | 12251 | 49 | 36 |
| 70% LMS | 5.18 | 2.55 | 10 | 6 | 3817 | 20 | 34 | 12592 | 49 | 31 |

**Additional Table S7.** Heavy metal content of leaves and tubers post-growth in various compost ratios. Heavy metals As, Cd, Cr, Co, and Pb were also analyzed but were below quantifiable levels (BQL) for all samples. Data is based on the analysis of pooled biological replicates for each treatment.

| Concentrations (mg kg^-1^) | | | | | | | | | | |
| --- | --- | --- | --- | --- | --- | --- | --- | --- | --- | --- |
|  | Al | | Cu | | Fe | | Ni | | Zn | |
| Soil Type | Leaf | Tuber | Leaf | Tuber | Leaf | Tuber | Leaf | Tuber | Leaf | Tuber |
| Adkins | 45 | 174 | 26 | 17 | 82 | 255 | 1 | BQL | 39 | 22 |
| 100% LMS | 37 | 198 | 180 | 37 | 118 | 255 | 5 | BQL | 53 | 31 |
| 85% LMS | 62 | 68 | 6 | 8 | 180 | 52 | 1 | BQL | 37 | 24 |
| 70%LMS | 46 | 45 | 5 | 4 | 122 | 54 | BQL | BQL | 32 | 19 |

**Additional Table S17.** Pre-growth assessments of the lunar regolith simulants used in Experiment 2. Data is based on the analysis of pooled biological replicated samples for each treatment.

|  | Moisture  (%) | pH | EC  (dS m^-1^) | Elemental Quantities  (mg kg^-1^) | | | | | | | | | Microbial respiration  (µg CO2-C/g dry soil/day) | | Organic matter (%) |
| --- | --- | --- | --- | --- | --- | --- | --- | --- | --- | --- | --- | --- | --- | --- | --- |
|  |  |  |  | Al | Ca | Cu | Fe | K | Mg | Mn | P | Zn | 24 hr | 96 hr |  |
| JSC | 11.03 | 7.50 | 0.54 | 1120.39 | 1968.11 | 2.32 | 1433.93 | 235.32 | 593.83 | 39.33 | 27.07 | 2.54 | 4.44 | 2.36 | 1.35 |
| LHS | 13.49 | 7.39 | 0.61 | 867.07 | 2052.82 | 1.36 | 440.12 | 92.84 | 209.96 | 12.48 | 24.74 | 1.25 | 6.84 | 3.34 | 2.23 |
| LMS | 1.00 | 7.61 | 0.43 | 810.62 | 1621.74 | 2.51 | 1104.08 | 152.56 | 466.92 | 30.65 | 39.60 | 1.81 | 9.82 | 5.04 | 1.45 |
| OPR | 10.99 | 7.72 | 0.60 | 1146.56 | 5848.36 | 2.33 | 3059.76 | 58.60 | 236.75 | 48.73 | 15.32 | 8.77 | 11.45 | 4.71 | 2.76 |
